# IFIT3 RNA-binding activity promotes influenza A virus infection and translation efficiency

**DOI:** 10.1101/2025.02.17.638785

**Authors:** Owen M. Sullivan, Daniel J. Nesbitt, Grace A. Schaack, Elizabeth Feltman, Thomas Nipper, Supasek Kongsomros, Sevilla G. Reed, Sarah L. Nelson, Cason R. King, Evgenia Shishkova, Joshua J. Coon, Andrew Mehle

**Affiliations:** Department of Medical Microbiology and Immunology, University of Wisconsin-Madison, USA; Department of Chemistry, University of Wisconsin-Madison, Madison, WI, USA; Department of Microbiology, Faculty of Science, Mahidol University, Thailand; Department of Biomolecular Chemistry, University of Wisconsin – Madison, Madison, WI 53506, USA; National Center for Quantitative Biology of Complex Systems, Madison, WI 53706, USA; Morgridge Institute for Research, Madison, WI 53515, USA

**Keywords:** Influenza A virus, IFIT3, innate immunity, RNA-binding, pro-viral, host interactions

## Abstract

Host cells produce a vast network of antiviral factors in response to viral infection. The interferon-induced proteins with tetratricopeptide repeats (IFITs) are important effectors of a broad-spectrum antiviral response. In contrast to their canonical roles, we previously identified IFIT2 and IFIT3 as pro-viral host factors during influenza A virus (IAV) infection. During IAV infection, IFIT2 binds and enhances translation of AU-rich cellular mRNAs, including many IFN-simulated gene products, establishing a model for its broad antiviral activity. But, IFIT2 also bound viral mRNAs and enhanced their translation resulting in increased viral replication. The ability of IFIT3 to bind RNA and whether this is important for its function was not known. Here we validate direct interactions between IFIT3 and RNA using electromobility shift assays (EMSAs). RNA-binding site identification (RBS-ID) experiments then identified an RNA-binding surface composed of residues conserved in IFIT3 orthologs and IFIT2 paralogs. Mutation of the RNA-binding site reduced the ability IFIT3 to promote IAV gene expression and translation efficiency when compared to wild type IFIT3. The functional units of IFIT2 and IFIT3 are homo- and heterodimers, however the RNA-binding surfaces are located near the dimerization interface. Using co-immunoprecipitation, we showed that mutations to these sites do not affect dimerization. Together, these data establish the link between IFIT3 RNA-binding and its ability to modulate translation of host and viral mRNAs during IAV infection.

**Importance:** Influenza A viruses (IAV) cause considerable morbidity and mortality through sporadic pandemics as well as annual epidemics. Zoonotic IAV strains pose an additional risk of spillover into a naive human population where prior immunity can have minimal effect. In this case, the first line of defense in the host is the innate immune response. Interferon stimulated genes (ISGs) produce a suite of proteins that are front-line effectors of innate immune responses. While ISGs are typically considered antiviral, new work has revealed an emerging trend where viruses co-opt ISGs for pro-viral function. Here, we determine how the ISG IFIT3 is used by IAV as a pro-viral factor, advancing our understanding of IFIT3 function generally as well as specifically in the context of IAV infection.

## Introduction

Influenza virus is a serious public health threat causing significant morbidity and mortality. Humans are under continual assault by seasonal influenza virus outbreaks, as well as sporadic pandemics with the potential for widespread infection and disease(1, 2). Pandemics are a deep concern for the human population as infection by these viruses are often associated with increased disease severity. Pandemic viruses emerge from animal reservoirs (3). Viruses with multiple animal reservoirs and rapid mutation rates pose a high risk for pandemic potential, and influenza A viruses (IAVs) fall into both of those categories. Spillover of IAV strains from zoonotic reservoirs have led to influenza pandemics in 1918, 1957, 1968 and 2009 (4). In addition to pandemics, IAVs cause annual epidemics with significant public health and economic burden. Vaccination and previous infections provide some protection against seasonal IAV infection (5). However, adaptive immune responses require prior exposure and are limited due to viral evolution selecting for immune escape variants during seasonal infections. In the absence of an effective adaptive immune response, the body’s first line of defense is innate immunity. The innate immune response requires no immune memory, which makes it even more essential in the case of pandemics where the human population has no prior virus exposure (6).

The type I interferon (IFN) response is the main signaling pathway for innate immune activation during viral infection. This pathway is initiated via host sensors that detect pathogen-associated molecular patterns (PAMPs) in infected cells (7). For a negative-sense RNA virus like IAV, retinoic acid inducible gene I (RIG-I) is the dominant sensor initiating innate immune responses. RIG-I senses viral RNA leading to the transcriptional upregulation of type I IFN and other first-wave effectors (8). Type I IFN then signals in an autocrine and paracrine fashion to initiate another transcriptional cascade that leads to expression of interferon stimulated genes (ISGs) (9). Conservative estimates suggest that ISGs consist of around 300 effector proteins with different effector functions that can interfere with every stage of the viral life cycle (10). The impact of these effector proteins is so pronounced that IAVs encode a protein, NS1, dedicated to antagonizing the type I IFN response. IAV lacking NS1 is severely impaired for viral replication and has significantly reduced pathology (11).

The interferon induced protein with tetratricopeptide repeats (IFIT) proteins are some of the first and most highly expressed ISGs. IFITs are a family of proteins characterized by their ability to bind RNA, and in some cases this RNA-binding function is connected to antiviral activity (12). There are five IFIT proteins in humans: IFIT1, IFIT1B, IFIT2, IFIT3, and IFIT5. IFIT1 recognizes and binds aberrant 5’ caps on mRNAs (13, 14). Most mRNAs obtain a specialized *N*7-methylguanosine (m7G) cap at their 5’ ends, creating the cap0 structure. The body of the message is also normally 2’O methylated on the first residue, creating the cap1 structure. The second residue can also be 2’O methylated, resulting in cap2 structures (15). IFIT1 specifically recognizes the incomplete cap0 structure, thereby inhibiting translation of these mRNAs (16). One example of this antiviral mechanism is seen in flaviviruses using a West Nile virus mutant that can no longer properly methylate viral mRNAs. IFIT1 binds the 5’ cap0 structures on viral transcripts leading to lower viral protein expression, reduced viral replication, and decreased pathogenicity in animals (17). IFITs can also exert antiviral activity by binding to host mRNAs. IFIT2 binds AU-rich sites in host mRNAs, especially ISG transcripts.

IFIT2 enhances the translation efficiency of the mRNAs it binds, leading to higher levels of ISG proteins and ultimately contributing to the antiviral state for the infected cell (18). IFIT5 also binds host RNAs such as tRNAs and RNA polymerase III transcripts. Limited reports suggest IFIT5 expression confers an antiviral phenotype, although it is less understood if RNA-binding is directly linked to this antiviral activity (19). Expression and knockdown studies have implicated IFIT3 as an antiviral protein. In humans, IFIT3 enhances the specificity and activity of IFIT1 (13, 14). However, whether IFIT3 binds RNAs, and if RNA binding is important for antiviral activity is unknown. Despite IFITs being broadly antiviral, in certain settings IFITs can be repurposed to promote replication. IFIT2 is co-opted by IAV where it binds viral mRNAs and enhances translation of viral proteins, shifting the balance from the canonical antiviral state to one that supports higher levels of viral replication (18).

Given the dual role played by IFIT2 during viral infection, we asked whether other IFIT proteins may also function to enhance IAV replication. IFIT3 emerged as a putative pro-viral factor for IAV in two different types of genetic screens (18, 20). Here we demonstrate that IFIT3 enhances IAV replication and define the molecular features of its activity. We show that IFIT3 promotes viral replication by increasing translation of viral proteins. IFIT3 heterodimerizes with IFIT2 as well as functions on its own but acts synergistically with IFIT2 to promote translation of viral mRNA. While IFITs play a central role in regulating replication of diverse viral infections, IAV exploits the antiviral activity of both IFIT2 and IFIT3 to enhance replication. We further show that IFIT3 is a *bona fide* RNA-binding protein that binds viral mRNAs. Using RNA-binding site identification (RBS-ID), we identified residues that create the RNA-binding site on IFIT3.

Mutagenesis of these residues showed that the RNA-binding sites are important for promoting viral gene expression. Thus, IFIT3 is an antiviral RNA-binding protein that is repurposed by IAV to promote replication.

## Results

### IFIT3 is pro-viral for influenza virus

The IFIT family has evolved through extensive gene duplication and gene conversion (21). IFIT2 and IFIT3 are orthologs with 57% identity. Although IFIT2 is typically considered an antiviral ISG, we previously showed that it stimulates expression of IAV proteins and promotes viral replication (18). We therefore tested if IFIT3 has similar functions during IAV infection. To determine if IFIT3 expression increases IAV replication, we performed transcriptional regulation by pathogen-programmed Cas9 (TRPPC) (20). This method utilizes sgRNAs encoded in the IAV genome to program CRISPR activation (CRISPRa). The virally encoded sgRNA direct catalytically inactive Cas9 and transcriptional activators to specific sites in the host genome, activating expression of the target gene. A549 lung cells stably expressing the CRISPRa proteins were infected with TRPPC virus encoding an sgRNA targeting IFIT3, or a non-targeting control (Figure 1A). The IFIT3-targeting virus increased IFIT3 mRNA levels by ∼15-fold compared to the control virus. This correlated with an increase in the levels of viral NP in these cells (Figure 1B). Infection with the IFIT3-targeting virus produced 5-10 fold more virus than the non-targeting control in both single-cycle and multi-cycle replication assays (Figure 1C-D). We then tested infection in cells lacking IFIT3 (Figure 1E). *IFIT3^-/-^* cells were infected with an influenza reporter virus that expresses nanoluciferase, enabling measures of viral gene expression. Viral gene expression was reduced in cells lacking IFIT3, similar to results from cells lacking IFIT2. Together, these data identify IFIT3 as another member of the IFIT family that enhances IAV replication.

**Figure 1:**
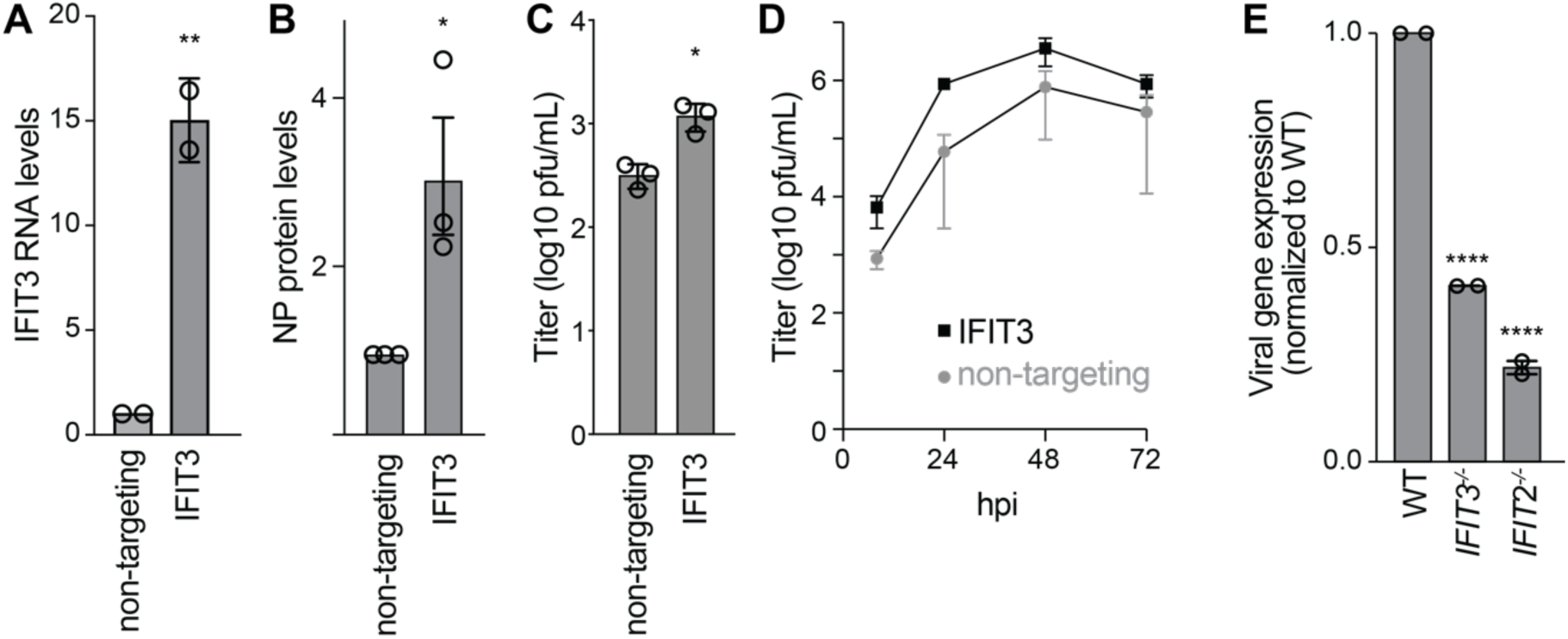
IFIT3 enhances influenza A virus replication. A) IFIT3 RNA expression measured by RT-qPCR from A549 CRISPRa cells inoculated with non-targeting or IFIT3-targeting TRPPC viruses (MOI = 5). Data are normalized to the non-targeting control. B) Influenza virus NP protein levels measured 8 hpi in A549 CRISPRa cells inoculated with non-targeting or IFIT3-targeting TRPPC viruses (MOI = 5). Data are normalized to the non-targeting control. C) Single-cycle replication was measured in A549 CRISPRa cells infected with virus targeting IFIT3 or a non-targeting control (MOI = 0.05). Viral titers were determined 18 hpi. D) Multicycle replication kinetics in A549 CRISPRa cells inoculated with virus targeting IFIT3 or a non-targeting control (MOI = 0.05). Titers were determined at the indicated time points by plaque assay. Data in A-D are mean of n=3 ± SD. Significance assessed by two-way Student’s t-test. E) Viral gene expression during single-cycle infection in WT, *IFIT3^-/-^* and *IFIT2^-/-^* A549 cells using a WSN-based reporter virus measured 8 hpi. Data are grand mean ± SEM of two independent biological replicates, each with 3 technical replicates. Significance was assessed by one-way ANOVA with Dunnett’s multiple comparisons test. * p < 0.5, ** p < 0.01, **** p < 0.0001.

IFIT3 and IFIT2 form homo- and heterodimers (13, 16, 22). We used our knockout cells to test the role of these different complexes. WT, *IFIT2^-/-^* and *IFIT3^-/-^* cells were transfected with plasmids to express IFIT2, IFIT3 or both IFIT2 and IFIT3 and subsequently infected with IAV. Pre-expressing IFIT2 or IFIT3 increased viral gene expression in WT and knockout cells, indicating that IFIT2 functions in the absence of IFIT3, and IFIT3 functions in the absence of IFIT2 (Figure 2A). However, we repeatedly detected the biggest enhancement of IAV gene expression when both IFIT2 and IFIT3 were pre-expressed. Higher levels of viral gene expression resulted in increased viral replication (Figure 2B). Similar patterns were observed, where co-expression of IFIT2 and IFIT3 resulted in the highest viral yields. Blotting shows that total IFIT levels did not dramatically increase when both IFIT2 and IFIT3 are co-expressed (Figure 2C), suggesting that the heterodimer is more active, consistent with it being the thermodynamically preferred complex (13). We note that while IFIT2 enhanced viral gene expression, it was routinely expressed at lower levels than IFIT3. In addition, as these complementation experiments restore activity, they show that phenotypes for our *IFIT2^-/-^* and *IFIT3^-/-^* cells are not due to background effects.

**Figure 2:**
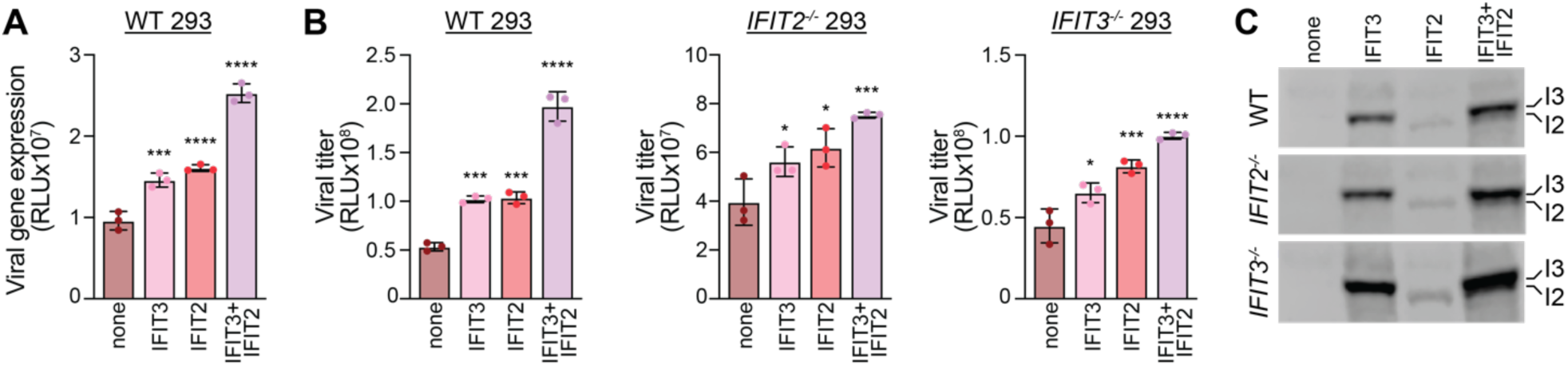
IFIT2 and IFIT3 function independently while also exhibiting additive pro-viral effect. A) WT 293 cells were transfected with plasmids expressing IFIT2, IFIT3, both IFIT2 and IFIT3, or an empty vector control. Cells were subsequently inoculated with a WSN-based PASTN reporter virus (MOI = 0.01) and viral gene expression was measured 8 hours post infection. B) Viral spread was measured in WT, *IFIT2^-/-^*and *IFIT3^-/-^* 293 cells following the same approach as in A, except infection was allowed to proceed for 24 hr before measuring reporter activity. C) Representative western blot of cell lysate before infection demonstrating IFIT2 and IFIT3 expression levels. Data are mean ± SD of a representative biological replicate from a total of 3. Significance assessed by one-way ANOVA with Dunnett’s multiple comparisons test. * p < 0.5, *** p < 0.001, **** p < 0.0001.

### IFIT3 binds host and viral RNA

Most IFIT family members have been characterized as RNA binding proteins (12), but whether IFIT3 binds RNA is unknown. IFIT3 interacts with IFIT2 and IFIT1, both of which are known to bind RNA. To directly assess the ability of IFIT3 to bind RNA on its own, we performed *in vitro* electromobility shift assay (EMSA). Purified recombinant IFIT3 was incubated with ∼200nt fragments of viral RNA, or a control RNA containing sequence from GFP. These specific segments of viral RNA were chosen as they are known IFIT2 binding sites during infection (18). IFIT3 specifically bound to 3 of the 4 viral RNAs, but only low levels of the GFP control (Figure 3A). IFIT3:RNA complexes assembled in a dose-dependent fashion (Figure 3B). The specificity of RNA binding activity was further demonstrated when complexes were lost upon boiling samples prior to electrophoresis (Figure 3C). This is the first formal demonstration that IFIT3 binds to RNA.

**Figure 3:**
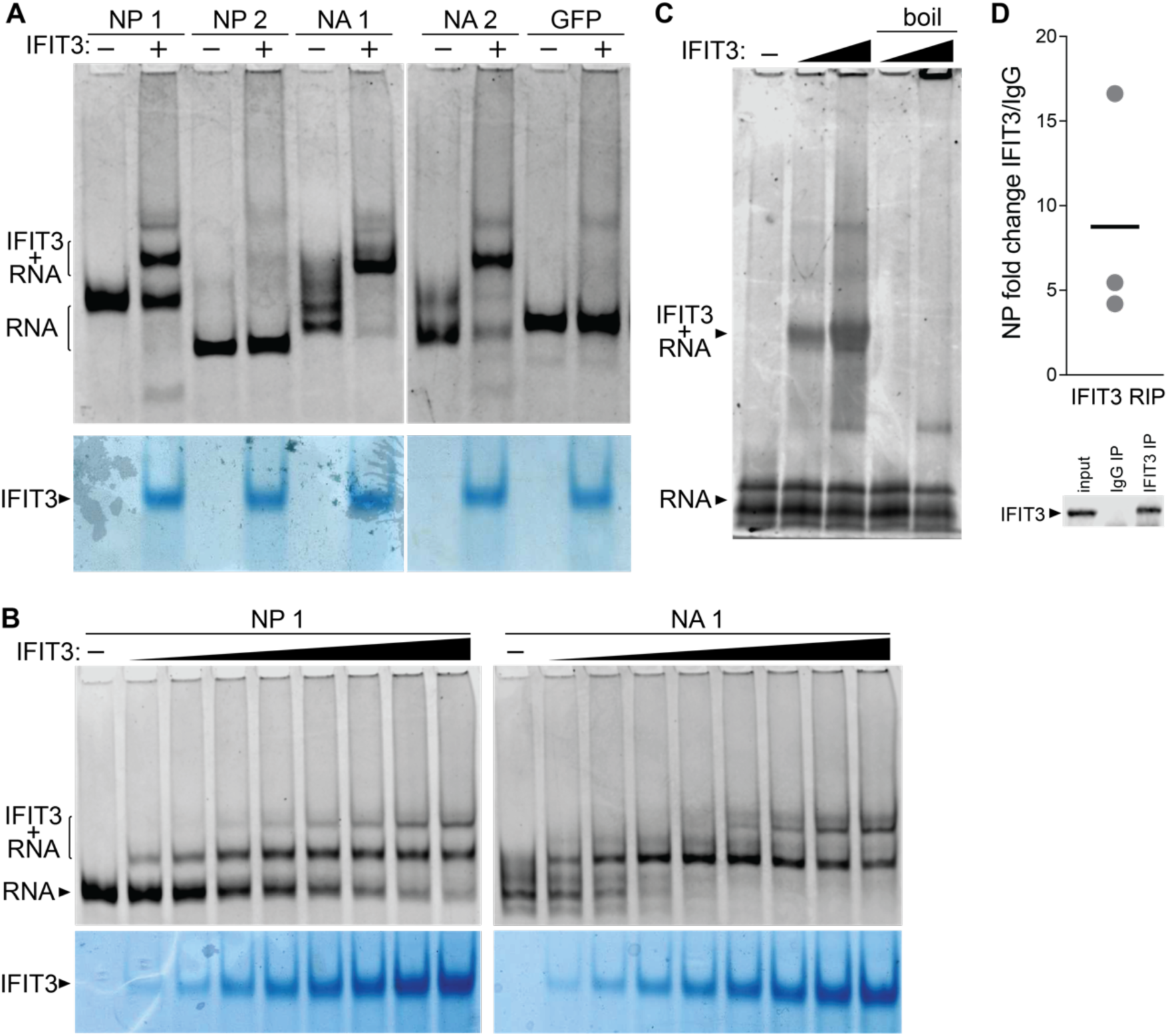
IFIT3 directly binds RNA. A) Electromobility shift assays (EMSA) were performed using *in vitro* transcribed RNAs derived from regions of the indicated viral genes in the presence or absence of recombinant IFIT3. RNAs was detected by Sybr gold staining (top) while proteins were visualized with Coomassie staining (bottom). B) EMSAs were performed with increasing amounts of recombinant IFIT3 and the indicated RNAs, revealing dose-dependent RNA binding. RNA was detected by Sybr gold staining (top) while protein was visualized with Coomassie staining (bottom). C) EMSAs were performed by forming complexes between NA1 RNA and increasing amounts of IFIT3. Where indicated, complexes were denatured by boiling before electrophoresis. D) RNA-immunoprecipitations (RIPs) were performed on infected cells lysates. *IFIT2^-/-^* A549 cells were infected with WSN (MOI = 0.02) for 24 h, lysed, and lysates were immunoprecipitated with an antibody targeting IFIT3 or an IgG control. IFIT3-specific binding was quantified by RT-qPCR relative to an IgG control. A representative Western blot showing expression and specific capture of IFIT3 is shown below. in the input as well as pulldown with the IFIT3 antibody, but not the IgG antibody.

To determine whether IFIT3 binds RNA *in vivo*, we performed RNA-immunoprecipitations from infected cells. To ensure RNA binding was mediated by IFIT3, and not a heterodimeric IFIT2:IFIT3 complex, we infected A549 *IFIT2^-/-^* with influenza A virus.

Infected cell lysates were immunoprecipitated with antibody against IFIT3 or an IgG control, total RNA was extracted, and samples were quantified by RT-qPCR (Figure 3D). IFIT3 enriched viral RNA 4-16 fold from the NP gene when compared to the IgG negative control. Together, these data demonstrate IFIT3 binds viral RNA under biologically relevant conditions.

### Identification of functionally important IFIT3 RNA binding sites

To determine whether RNA-binding contributes to IFIT3 activity, we first needed to identify what region of IFIT3 binds to RNA. We therefore performed RNA-binding site identification (RBS-ID) for IFIT3 (23). For RBS-ID, cells were UV crosslinked to covalently capture RNA:protein interactions, IFIT3 was purified from cell lysates, RNA and protein were digested, and mass spectrometry was performed to identify the specific residues that crosslinked to RNA. We expressed and purified IFIT3 from cells that were treated with ifNβ to mimic an antiviral response. RBS-ID revealed 3 high-confidence RNA:protein crosslink sites in IFIT3: C239, C283, and S360/H361 (Figure 4A-D). The exact crosslink site at S360/H361 could not be resolved.

**Fig. 4:**
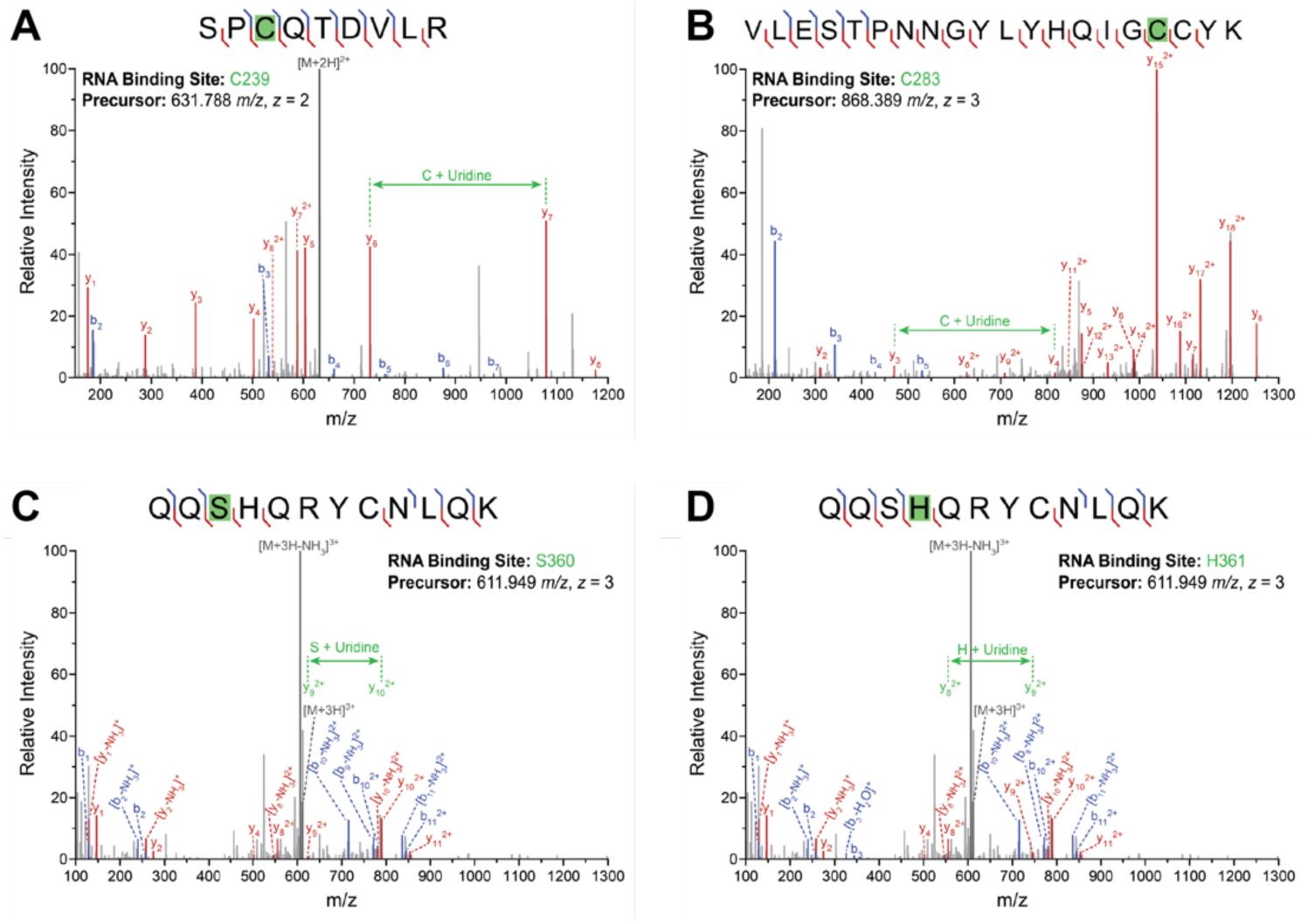
RNA-binding site identification on IFIT3. A-D) A-D) RBS-ID was performed on cross-linked IFIT3:RNA complexes purified from IFN-treated cells. Mass spectra highlight modified peptides and the specific residue cross-linked to the bound RNAs. Red and blue annotations denote y- and b-type sequence fragments, respectively, and the peptide precursor is denoted in gray. Localization of the cross-linking modification is shown in green through the mass difference between fragments of the same type adjacent to the modification site.

Residues C239, C283 and H361 are highly conserved across IFIT3 orthologs from diverse influenza virus hosts and animal models (Figure 5A). IFIT3 C283 and H361 sites are also conserved at analogous positions in human IFIT2. Full-length IFIT3 structures have not yet been reported, therefore, to map RNA binding sites we predicted its structure using the IFIT2 homodimer structure as a model (Figure 5B). The C-terminus of each IFIT2 monomer forms a super helical structure with a large positively charged surface on the interior facing the other monomer that has been shown to be important for RNA binding (24). The IFIT3 model also predicts a similar positively charged surface, although not as extensive as IFIT2. The predicted basic patch in IFIT3 contains analogous residues that when mutated in IFIT2 disrupt RNA binding; IFIT2 K255, R259, and R292 map to IFIT3 K249, R252/R253 and K286, respectively (outline in Figure 5A) (24). Like the RBS-ID sites, the basic residues are highly conserved in IFIT3 orthologs (Figure 5A). Residues identified by RBS-ID flank the predicted basic patch, with C283 adjacent to it and S360/H361 across the helical coil.

**Figure 5:**
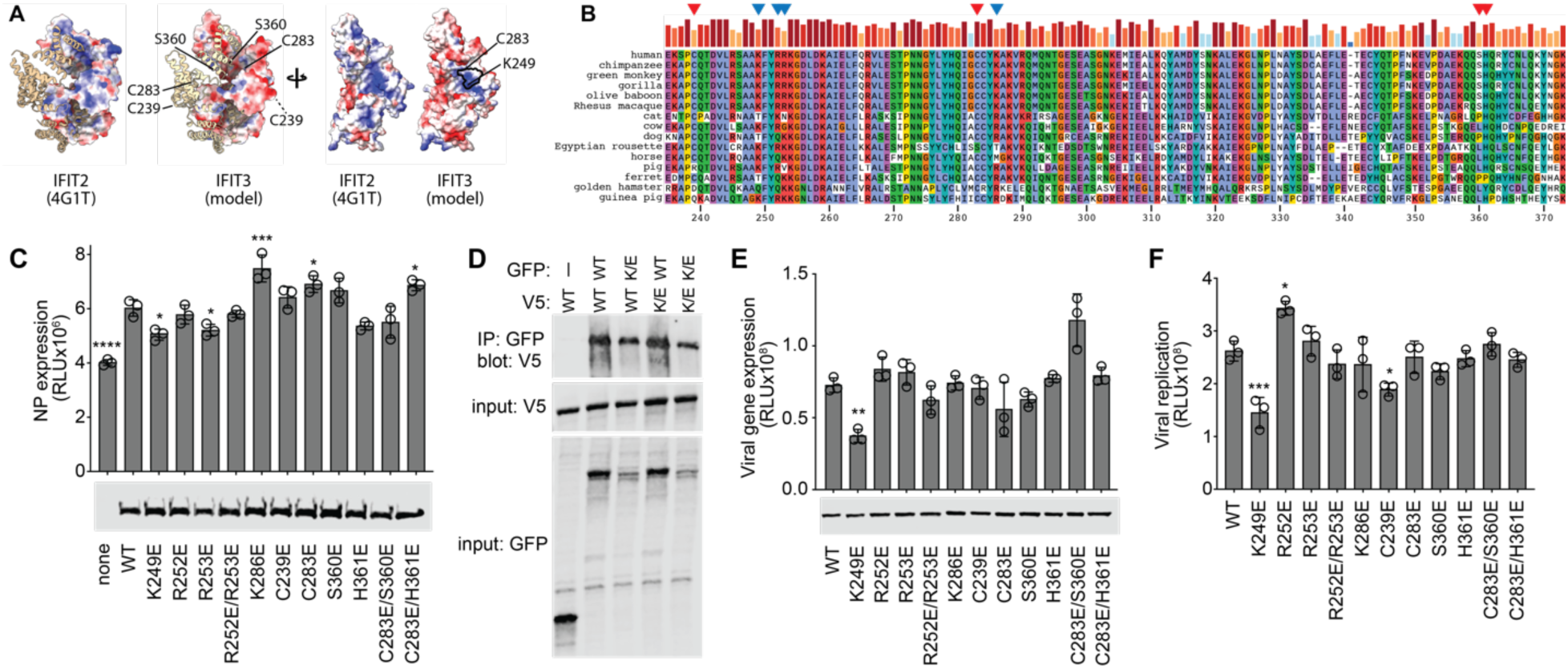
R**N**A **binding by IFIT3 is important for its pro-viral function** A) A putative IFIT3 structure was modeled on the IFIT2 structure. Dimers (left) are depicted as pairs of ribbon and space-filling subunits, while monomers (right) are shown rotated to emphasize surfaces facing the dimer interface. The computed electrostatic surface potential is shown from blue (basic) to red (acidic). Residues identified by RBS-ID as cross-linking to RNA are indicated and a conserved basic patch is outlined in black. B) Protein alignment of a portion of IFIT3 from multiple influenza A virus host. Red arrows indicate sites identified via RBS-ID and blue arrows indicate amino acids that were mutated based on sequence similarity to IFIT2 RNA-binding mutants. C) Influenza NP expression reporter assays were performed in *IFIT3^-/-^* 293 cells co-transfected WT or mutant IFIT3 or an empty-vector control (above). Equivalent expression of WT and mutant IFIT3 was shown by Western blot. D) IFIT3 dimer formation was measured for WT and K249E (K/E) IFIT3. WT and mutant IFIT3 were expressed as GFP- and V5-tagged proteins in *IFIT3^-/-^* 293 cells, the GFP-tagged form was immunoprecipitated from cell lysates, and co-precipitating proteins were detected by blotting for V5-tagged versions. Co-precipitating proteins are shown in the top blot, while input proteins are detected in the middle and bottom blots. E) *IFIT3^-/-^*293 cells were transfected with plasmids expressing WT or mutant IFIT3 and subsequently inoculated with a WSN-based PASTN reporter virus (MOI = 0.01). Viral gene expression was measured 8 hours post infection. IFIT3 expression was detected by western blot. F) Viral spread was measured in *IFIT3^-/-^*293 cells following the same approach as in E, except infection was allowed to proceed for 24 hr before measuring reporter activity. Data are mean ± SD of 3 technical replicates from a representative biological replicate from a total of 3 biological replicates. Significance assessed by one-way ANOVA with Dunnett’s multiple comparisons test. * p < 0.5, ** p < 0.01, *** p < 0.001 **** p < 0.0001.

We tested the importance of RNA binding in a series of IFIT3 activity assays. We had previously shown that IFIT2 stimulates translation of viral mRNAs leading us to predict that IFIT3 may serve the same function (18). We tested this in a reporter assay where the open reading frame of NP is fused to luciferase, allowing us to quantify viral protein levels. Assays were performed in 293 *IFIT3^-/-^* cells to remove the contribution of endogenous IFIT3. Expression of WT IFIT3 stimulated production of viral NP (Figure 5C). Specific mutations that disrupt the predicted basic patch eliminated IFIT3 activity, notably K249E. Interestingly, none of the residues identified by RBS-ID were essential for IFIT3 function, suggesting that while they are proximal to the bound RNA they are not essential for RNA binding. All IFIT3 proteins expressed at similar levels.

The most stable form of IFIT3 (and IFIT2) is a dimer. The RNA-binding surface we defined abuts the dimer interface. To ensure the above phenotypes were not due to inadvertently disrupting dimerization, we tested IFIT3:IFIT3 interactions between WT and RNA-binding mutants. GFP- and V5-tagged versions of IFIT3 and IFIT3 K249E were co-expressed in cells. Cells were lysed, IFIT3 complexes were immunopurified via the GFP tag, and co-precipitating IFIT3 was detected via the V5 tag. IFIT3 was co-precipitated by IFIT3-GFP, but not the GFP-only control, demonstrating specific interaction (Figure 5D). IFIT3 K249E was also efficiently co-precipitated by IFIT3-GFP, even though it is expressed at lower levels than WT. Reciprocal interactions were also detected, in addition to formation of IFIT3 K249E dimers. Thus, IFIT3 K249E retained its ability to form dimers.

Finally, we tested the importance of IFIT3 RNA-binding activity during influenza A virus infection. We again used *IFIT3^-/-^* cells, expressing the desired IFIT3 via transfection prior to infection. Compared to cells expressing WT IFIT3, viral gene expression was significantly reduced during the early stages of infection in cells expressing IFIT3 K249E (Figure 5E) and this correlated with a drop in viral spread at later times of infection (Figure 5F). All IFIT3 proteins were expressed at similar levels. Combined, these data identify IFIT3 K249 as a part of the RNA-binding region identified by RBS-ID and a key residue that is important for IFIT3 activity.

## Discussion

Influenza virus poses a significant threat to human health, especially in the case of pandemic influenza viruses (1). Typically, influenza pandemics are spurred by exposure of immunologically naïve populations to novel influenza virus strains. Without immune memory from previous infections or vaccines, infected hosts rely on the innate immune response to slow infection until an adaptive immune response can be mounted (6). However, recent work has demonstrated that viruses can usurp innate immune effectors to enhance viral replication (25). Here we show that IFIT3, one of the first and most abundantly expressed innate immune effectors, is repurposed by IAV to support its replication. IFIT3 promoted influenza virus replication on its own and worked additively with the family member and binding partner IFIT2. Orthogonal experimental approaches revealed that IFIT3 is an RNA-binding protein and identified a conserved basic patch that forms the RNA-binding site. Mutating the conserved basic patch disrupted the ability of IFIT3 to enhance IAV gene expression and replication, connecting RNA binding to IFIT3 activity. These data reveal mechanisms for the pro-viral activity of IFIT3 and shed light on how it likely functions in its canonical antiviral role.

The IFITs are important effectors of the innate immune system suppressing replication of a diverse array of viruses through multiple mechanisms (12). Our work here and elsewhere defined a counterexample where IFIT2 and IFIT3 have pro-viral roles during influenza virus infection. IFIT2 binds to and enhances translation of IAV transcripts during infection to increase viral protein production, and ultimately viral replication (18). We now show that IFIT3 also binds RNA and stimulates expression of viral gene products. While the use of knockout cells showed that IFIT2 and IFIT3 can function independently, viral replication was highest when both IFIT2 and IFIT3 were expressed. IFITs have discrete functions depending on their binding partners (13, 14). The IFIT2/IFIT3 heterodimer is the thermodynamically preferred complex, reflected in the increased activity detected when both proteins were expressed (13). IFIT2 homodimers are known to enhance the apoptosis pathway, which has been implicated in its role in controlling certain types of cancers (12, 22, 26, 27). While apoptosis can be beneficial during influenza virus infection (28, 29), the apoptosis pathway is not involved in the pro-viral function of IFIT2 during IAV infection (18). IFIT3 is thought to redirect IFIT2 from the apoptosis pathway towards its translation effector function (12, 18, 30). This could help explain the additive effects of IFIT2 and IFIT3 reported here.

IFIT2 and IFIT3 enhance translation of target proteins, but the molecular mechanisms by which they achieve this function remains unclear. Previous work showed that the RNA-binding activity of IFIT2 is essential to pro-viral function (18). Our data now reveal that IFIT3 is also an RNA-binding protein. RNA binding was mapped to a basic patch that is shared with IFIT3 homologous in other species. An analogous region of IFIT2 is also important for its ability to bind RNA (24). Mutations to the RNA-binding patch in IFIT3 disrupted its ability to stimulate viral gene expression or replication but did not affect its ability to form homodimers. Thus, all members of the IFIT family have now been shown to bind RNA and this RNA-binding activity is central to their function (12). Whether IFIT3 binds to host mRNAs like IFIT2, and if the repertoire of bound RNAs differs between IFIT2, IFIT3 or IFIT2/IFIT3 dimers, remains to be determined.

IFIT2 binds to host ISG where to enhance their translation and establish a generic antiviral state (18). It is this same process that is exploited by influenza virus to promote translation of its own proteins. Likewise, we speculate that the properties defined here underlying the pro-viral function of IFIT3 during influenza virus infection are the same features that allow it to exert antiviral activities during other viral infection. Blindly targeting pro-viral proteins, without knowledge of their mechanism, could have unwanted and deleterious consequence. This highlights the importance of understanding the mode of action for host pro-viral factors.

## Acknowledgements

We thank members of the Mehle lab for feedback and critical reading of the manuscript and Nancy Reich and Nicholas Heaton for sharing reagents. This work was supported by National Institutes of Health grants R01AI164690 (AM), P41 GM108538 (JC), R35 GM118110 (JC), T32AI055397 (OMS), T32HG002760 (DN), T32AI007414 (TN), and AI007414 and GM140935 (GAS). CRK was an Open Philanthropy Fellow of the Life Sciences Research Foundation. AM was a Burroughs Wellcome Fund Investigator in the Pathogenesis of Infectious Disease and is currently an H.I. Romnes Faculty Fellow funded by the Wisconsin Alumni Research Foundation (USA) and a Vilas Mid-Career Investigator.

## Author Contributions

Conceptualization: OMS, GAS, AM

Methodology: OMS, DJN, GAS, EF, TN, SGR, SLN, CRK, SK, ES, AM

Formal Analysis: OMS, DJN, GAS, AM

Investigation: OMS, DJN, GAS, EF, TN, SGR, SLN, CRK, SK, ES, AM

Writing – Original Draft: OMS and AM Writing – Review & Editing: all Visualization: OMS, DJN, GAS, EF, TN, AM

Funding Acquisition: OMS, DJN, GAS, AM, and JJC Supervision: AM and JJC

## Methods

### Cells and cell culture

Mammalian cells were grown in DMEM supplemented with 10% FBS and maintained at 37°C, 5% CO_2_. Cells were routinely tested for mycoplasma (MycoAlert, Lonza). Human lung alveolar epithelial A549 cells (CCL-185), Madin–Darby canine kidney (MDCK) cells (CCL-34), human embryonic kidney 293T (HEK293T) cells (CRL-3216), and human embryonic kidney 293 (HEK293) cells (CRL-1573) were acquired from the American Type Culture Collection (ATCC). MDCK-SIAT1-TMPRSS2 cells were kindly shared by Jesse Bloom (31). A549 *IFIT3^-/-^*, A549 *IFIT2^-/-^*, 293 *IFIT3^-/-^*, and 293 *IFIT2^-/-^*cells were previously generated by CRISPR/Cas9 mutagenesis (18) via lentivirus transduction (A549) or produced by Synthego (293s), isolated as clonal lines, and genotyped. A549-SAM cells were generously provided by Nicholas Heaton (32).

### Plasmids

Plasmids encoding wild type human IFIT2 and IFIT3 with C-terminal V5 epitope tags or GFP fusions were a kind gift of N. Reich (Stony Brook University) (22). IFIT3 RNA-binding mutants were generated by inverse PCR and confirmed by sequencing. eGFP-IFIT3 K249E expression plasmid was generated using HiFi DNA assembly (NEB) of PCR products.

The IFIT reporter construct pcDNA3-NP-2A-Nluc was created using Gibson assembly to extend the open reading of NP with in-frame coding sequence for the porcine teschovirus-1 2A peptide and Nanoluciferase. For bacterial expression, IFIT3 was cloned into pET15b to acquire an N-terminal His tag. All plasmids were sequence verified.

### Viruses and virus production

Viral infections were performed using reporter viruses encoding the NanoLuc (Nluc) reporter gene as a polyprotein with PA, which have been previously described for the WSN (PASTN) and A/California/04/2009 (H1N1) strains (CA04-PASTN) (33, 34). TRPPCa IFIT3 virus was generated on an influenza virus A/PR8/34 (H1N1; PR8) backbone and plaque purified prior to amplification of stocks (20). Viruses were propagated as before in MDCK cells or MDCK-SIAT-TMPRSS2 cells in virus growth media (VGM) (OptiMEM, 1% penicillin/streptomycin, 2.5% HEPES, 0.2% BSA, 100μg/ml CaCl_2_) (35, 36). Virus growth media for MDCK cells was also supplemented with up to 2μg/ml TPCK-trypsin. Viral stocks were titered in triplicate by plaque assay. Cell culture infections were performed in triplicate at 37°C and inoculated at the indicated MOI.

### qRT-PCR

Total RNA was extracted from cells using TRIzol and mRNAs were reverse transcribed using an oligo-dT primer and MMLV reverse transcriptase. cDNA was diluted 1:10 in water, and qPCR was performed in technical duplicate with iTaq SYBR master mix (Biorad) in a Step One Plus RT-PCR instrument (Applied Biosystems). The following primers were used:

NP forward AGACCAGAAGATGTGTCTTTCC NP reverse CGCACCAGATCGTTCGAGTCG

Ct values of target genes were normalized to β-actin and relative gene expression levels between conditions and were calculated via the ΔΔCt method. Expression levels are plotted as fold-change over a baseline condition set to 1. Values are the mean of 3 biological replicates.

### TRPPC infections and reporters

TRPPC infections followed our prior approach (20). A549-SAM were seeded at 1.5×10^6^ cells per well in a 6-well plate and allowed to grow for 24 hours. Cells were then infected with TRPPC viruses at an MOI of 0.05 in VGM supplemented with 1.5 μg/mL of TPCK. After 1 hour, inoculum was removed and replaced with fresh VGM. Single-cycle replication was measured by harvesting virus 8 hpi. Multicycle infections continued for 72 hours with viral aliquots being taken at 12, 24, 48, and 72 hpi. Viral titers were determined for each aliquot by plaque assay on MDCK-SIAT-TMPRSS2 cells.

### Viral gene expression and replication

Nanoluciferase activity was measured as a proxy for viral gene expression (8 hpi) and viral replication (24 hpi) using PASTN (37). 293 cells (WT, *IFIT2^-/-^*, or *IFIT3^-/-^*) were inoculated with a nanoluciferase reporter virus at the indicated MOI. The inoculum was removed after 1 h and replaced with fresh viral growth media. For infection experiments testing IFIT complementation and IFIT3 mutants, 0.75 x 10^6^ 293 cells per well were plated in a 6-well plate for 24 h and transfected with the indicated DNA construct. 24 hours after transfection, cells were trypsinized and used to seed a white bottom 96-well plate for infection assay (3 wells per replicate) or a 24-well plate to measure protein expression. 24 hours after seeding, cells were infected as described above. Nanoluciferase activity was measured either 8 hpi or 24 hpi by incubating cells in 50 μL 1X Glo buffer (Promega) for 10 minutes at room temperature in order to lyse cells. NanoGlo reagent was added and analyzed based on manufacturer instructions (Promega).

### Translation reporter assay

The impact of IFIT3 on translation was measured using the NP-2A-Nluc fusion proteins. 293 cells were transfected with constructs expressing either NP-2A-Nluc in the presence or absence of wild type or mutant IFIT3. Nanoluciferase activity was measured by lysing cells in 50 mM Tris pH 7.4, 150 mM NaCl and 0.5% NP40 and analyzing an aliquot with the NanoGlo luciferase assay following the manufacturer’s instructions (Promega). Fold induction was calculated relative to the negative controls lacking IFIT3 and averaged across all replicates.

### Electromobility shift assay

His-TEV IFIT3 was expressed and purified from Rossetta2(DE3) bacteria (Novagen) following prior approaches (18). Briefly, proteins expressed in *E. coli* at 16 °C overnight and cells were lysed by sonication in high salt buffer (20 mM HEPES pH 7.4, 1 M NaCl) supplemented with 0.5 mg/ml lysozyme and 0.2 mM PMSF. Lysates were clarified by centrifugation and protein was purified in bulk using Ni-NTA Superflow (Qiagen) followed by ion exchange on a 1ml HiTrap Q column (GE Life Sciences). Purified protein was dialyzed against an excess of low salt buffer containing 10% glycerol. Proteins were quantified by Bradford assay, assessed by gel electrophoresis, aliquoted, flash frozen in liquid nitrogen and stored at −80 °C until use.

RNAs were transcribed with T7 RNA polymerase. Templates were made by PCR using a 5’ primer that appended the T7 promoter sequence onto the product to create NP 1-200, NP 1241-1440, NA 50-249, NA 802-1011 and eGFP 18-217 (numbering is nucleotide position in cRNA). RNA was purified by TRIzol extraction, quantified and verified by formaldehyde denaturing agarose gel electrophoresis.

Protein:RNA binding reactions were performed by assembling 20 μl reactions containing ∼25 nM mRNA and 0 or 12.5 μM protein in 20 mM HEPES, 150 mM NaCl, 2.5 mM MgCl_2_, 1 mM DTT and 4U RNasin. Binding was performed at room temperature for 45 min. Samples were separated on a 6% polyacrylamide gel buffered with 0.5x TBE 4 °C. After separation, the gel was stained with Sybr Gold and RNA was visualized.

### IFIT3 RIP qRT-PCR

*IFIT2^-/-^* A549 cells were infected with WSN (MOI = 0.02) for 24 h. Cells were lysed in co-immunoprecipitation buffer (50 mM Tris pH 7.5, 150 mM NaCl, 0.5% NP40, 1x protease inhibitor cocktail (Sigma), and 20U/mL RNasin (Promega)) and clarified. Lysates were incubated for 16 h with 3 μg of IFIT3 antibody or rabbit IgG (Cell Signaling Technologies 2729S) in the presence of 1mg/mL BSA. Immuno-complexes were captured with protein A agarose resin (Sigma) for three hours and washed 6 times with co-immunoprecipitation buffer. 10% of the resin was reserved for western blot analysis and RNA was extracted from the remainder with TRIzol, precipitated, and resuspended in 10 μl of water. Equal volumes of IFIT3-bound RNA and IgG-bound RNA were used for qRT-PCR analysis. RT-qPCR was performed as above. IFIT3-bound samples were normalized to IgG-bound samples using the ΔCt method.

### RBS-ID cell preparation

Cell preparation for RNA-binding site identification (RBS-ID) utilized 293T cells. 4×10^6^ cells were plated in 10 cm^2^ dishes in duplicate. 24 hours after seeding, cells were treated with 500U of recombinant ifNβ and transfected with DNA containing IFIT3-V5. 24 hours post transfection, cells were crosslinked using UV radiation to preserve RNA:protein complexes. Media was aspirated from cells and replaced with 1 mL of cold 1X DPBS. Cells were placed on ice and subject to UV cross-linking with 254 nm light at 400 mJ/cm^2^ followed by 200 mJ/cm^2^.

Crosslinked cells were washed 3 times with 1X DPBS. Duplicate dishes were pooled and cell pellets were flash frozen in liquid nitrogen before storing in −80°C until preparation.

Cell pellets were thawed on ice, lysed in RIPA buffer [100mM Tris Cl (pH 7.5), 150mM NaCl, 0.2% (w/v) SDS, 1% (w/v) DOC, 2% (v/v) Igepal CA-630] and clarified by centrifugation. IFIT3-V5:RNA complexes were purified by incubating lysate with magnetic anti-V5 nanobody trap resin (Proteintech) overnight with rocking at 4°C. Resin was then washed 4 times in RIPA, followed by 3 washed in 1X DPBS. RNA:protein complexes were then eluted with 35 μL of 8M urea for 30 minutes at 45°C, and 15% of the elution was run on SDS-PAGE and Coomassie stained for protein verification.

### Mass spectrometry protein digestion

First, the two eluates from each sample were pooled. For protein reduction and alkylation, a 10x solution of tris(2-carboxyethyl)phosphine (Sigma-Aldrich) and chloroacetamide (Sigma-Aldrich) was added to each sample (final concentrations 10 mM and 40 mM, respectively), and samples were shaken at ambient temperature for 15 minutes. Next, Lysyl Endopeptidase (Fujifilm Wako Chemicals) was added to each sample (1:50 enzyme:protein ratio by mass), and the samples incubated for 4 hours at ambient temperature. Samples were then diluted to a final urea concentration of 2M with 100 mM Tris pH 8.0 (Thermo-Fisher Scientific), then trypsin (Promega) was added to each sample (1:50 enzyme:protein ratio by mass), and the samples incubated for 4 hours at ambient temperature. Following digestion, samples were acidified with 10% trifluoroacetic acid (TFA, Thermo-Fisher Scientific) to a pH of ∼1.5.

C18 Sep-Pak desalting cartridges (50mg sorbent, Waters) were used for peptide desalting. Cartridges were first conditioned with 3 mL of acetonitrile (ACN, Thermo-Fisher Scientific, LC-MS grade); 1 mL of 70% ACN, 0.1% TFA in water; 1 mL of 40% ACN, 0.1% TFA in water; and 3 mL of 0.1% TFA in water. Samples were added then washed 3 mL of 0.1% TFA in water, then eluted with 750 µL of 40% ACN, 0.1% TFA in water and 750 µL of 70% ACN, 0.1% TFA in water and the eluates pooled. Peptides were then dried in a vacuum concentrator (SPD130DLX, Thermo-Fisher Scientific).

### Mass spectrometry hydrofluoric acid digestion

Dried peptides were reconstituted in 100 µL of 50 mM Tris, pH 8.0 in water. Next, 400 µL of concentrated hydrofluoric acid (HF, 48% w/w in water, Sigma-Aldrich) were added to each sample. Sample tubes were sealed and placed in an air-tight box with free calcium carbonate powder (Sigma-Aldrich) and incubated overnight at 4 °C. Samples were then dried in a vacuum concentrator (SPD131DDA, Thermo-Fisher Scientific) housed within a fume hood, connected to a plastic solvent collection trap filled with calcium carbonate and cooled with dry ice. Additional sample tubes were filled with calcium carbonate and sealed with an air-permeable membrane (Diversified Biotech) and placed within the vacuum concentrator. The dried samples were reconstituted with 500 µL of 0.1% TFA in water and desalted in the manner described in the previous section. Following desalting and vacuum-drying, samples were reconstituted in 0.2% formic acid (Thermo-Fisher Scientific, LC-MS grade), and the peptide concentration for each sample was measured by NanoDrop (Thermo-Fisher Scientific) using the absorbance at 205 nm.

### LC-MS data collection

Peptides were separated using a Neo Vanquish UHPLC System (Thermo-Fisher Scientific) on a 40 cm bare fused silica capillary (75 µm i.d., 360 µm o.d.) packed in-house (38) with 1.7 µm, 130 Å pore size BEH C18 particles (Waters) held at 55 °C with a custom-built column heater.

Solvents A and B consisted of 0.1% formic acid in water and 0.1% formic acid in 80:20 ACN:water, respectively. The flow rate was held constant at 300 nL/min, and 700 µg of peptides per sample were injected on column and separated with the following gradient: 0-1 mins, 0-6% B; 1-51 mins, 6-50% B; 51-52 mins, 50-100% B; 52-60 mins, 100% B.

Initial mass spectrometry data was collected using a quadrupole-Orbitrap-quadrupole linear ion trap Tribrid instrument (Orbitrap Ascend, Thermo-Fisher Scientific). The spray voltage was held constant at 2 kV in positive ion mode. MS1 scans were acquired in the Orbitrap in profile mode with the following parameters: resolving power, 60,000 at 200 m/z; scan range, 300-1350 m/z; maximum inject time, 123 ms; AGC target, 1,000,000(normalized AGC target, 250%).

Monoisotopic precursor selection was enabled in peptide mode, and precursors of charge states 2-6 were selected for MS2 fragmentation. Dynamic exclusion was was set to 15 seconds with a 25 ppm mass tolerance. Data-dependent MS2 scans were acquired in centroid mode throughout a 1.5 s cycle time with the following parameters: activation type, HCD; collision energy mode, fixed; normalized collision energy, 30%; resolving power, 15,000 at 200 m/z; scan range, 150-1350 m/z; maximum inject time, 85 ms; AGC target, 125,000; normalized AGC target, 250%.

To validate potential hits, time-scheduled PRM analysis of selected m/z and z targets was conducted (39). For PRM analysis, HCD MS2 scans were collected using stepped collision energies of 23%, 27%, and 32% (normalized) at an AGC target of 50,000 (100%) for each collision energy with a maximum inject time of 125 ms. The resolving power was set to 15,000 at 200 m/z and the scan range was 100-1350 m/z. The same amount of peptides was injected on column and separated using the same gradient parameters for each run.

### LC-MS data analysis

Initial LC-MS data were analyzed using the FragPipe (40, 41) software suite (v. 22.0). Spectra were searched with a Uniprot human database (reviewed sequences only, isoforms included, downloaded 29 Jan. 2024) with the modified IFIT3 sequence added. The default closed search settings were employed, but with some alterations. Up to 3 missed cleavages were permitted. Alkylation of cysteine was set as a fixed modification and methionine oxidation was set as a variable modification with one max occurrence. For the RBS-ID modifications, a variable modification of mass delta 187.04807 (uridine crosslink minus carbamidomethylation, C_7_H_9_NO_5_) was enabled on cysteine residues, with up to two max occurrences (23). For all non-cysteine residues, a variable modification of mass delta 244.06953 (uridine, C_9_H_12_N_2_O_6_) was enabled with up to two max occurrences (23). In total, up to three variable modifications per peptide were allowed. From the closed search, peptide spectral matches with crosslink modifications were manually inspected and targeted for subsequent PRM experiments. From the PRM data, spectral data for each peptide were extracted from Thermo Scientific Freestyle, annotated with Interactive Peptide Spectral Annotator (IPSA) (42) and manually validated.

### IFIT3 protein alignment and modeling

A predicted structure for IFIT3 was created by using a monomer of IFIT2 (PDB 4G1T) as a template in SWISS-MODEL (24, 43). Conserved residues were identified by using CLUSTAL to align human IFIT3 to orthologs from other mammals.

### Western Blot

Western blots were performed on clarified cell lysates that were lysed in 50 mM Tris pH 7.4, 150 mM NaCl, and 0.5% NP40. Samples were denatured and transferred to nitrocellulose membranes and blocked in a 1X solution of BSA. Membranes were then probed with primary antibodies targeting V5 (Bethyl Laboratories, Inc., cat. no. A190-220A) at 1:5000 dilution, IFIT3 (Proteintech 15201-1-AP) at 1:800 dilution, IFIT2 (Proteintech 12604-1-AP), at 1:500 dilution or GFP (Proteintech 50430-2-AP) at a 1:1000 dilution. Membranes were then incubated with secondary antibodies compatible with IR imaging at 1:10000 dilutions. Antigens were detected via IR using and Odessey Fc Imager equipped with Image Studio (LI-COR).

### Co-immunoprecipitations

Interactions between wild type and mutant IFIT3 were tested by co-immunoprecipitation. WT 293T cells were transfected with combinations of wild type and mutant IFIT3-V5 or IFIT3-eGFP. Cells were then lysed in co-IP buffer and clarified by centrifugation. Lysates were immunoprecipitated overnight with GFP trap resin (Proteintech gta). Resin was washed 4 times in co-IP buffer and proteins were eluted by boiling in 1x Laemmli loading buffer. Eluted proteins were detected by western blot.

### Statistical Analysis

Assays were performed with three technical replicates and represent at least three independent biological replicates. Data are presented as mean and SD for technical replicates or grand mean and SEM of independent biological replicates. When data were normalized, error was propagated to each individual experimental condition. Statistical significance was determined for pairwise comparisons with a Student’s t test and for multiple comparisons an ANOVA and *post hoc* Dunnett’s or Sidak’s test. Statistical tests were performed using Prism (GraphPad).

